# Modeling pupil responses to rapid sequential events

**DOI:** 10.1101/655902

**Authors:** Rachel N. Denison, Jacob A. Parker, Marisa Carrasco

**Affiliations:** Department of Psychology and Center for Neural Science, New York University, New York, NY 10003, USA; Human Motor Control Section, Medical Neurology Branch, National Institute of Neurological Disorders and Stroke, NIH, Bethesda, MD, USA

## Abstract

Pupil size is an easily accessible, noninvasive online indicator of various perceptual and cognitive processes. Pupil measurements have the potential to reveal continuous processing dynamics throughout an experimental trial, including anticipatory responses. However, the relatively sluggish (∼2 s) response dynamics of pupil dilation make it challenging to connect changes in pupil size to events occurring close together in time. Researchers have used models to link changes in pupil size to specific trial events, but such methods have not been systematically evaluated. Here we developed and evaluated a general linear model (GLM) pipeline that estimates pupillary responses to multiple rapid events within an experimental trial. We evaluated the modeling approach using a sample dataset in which multiple sequential stimuli were presented within 2-s trials. We found: (1) Model fits improved when the pupil impulse response function (puRF) was fit for each observer. PuRFs varied substantially across individuals but were consistent for each individual. (2) Model fits also improved when pupil responses were not assumed to occur simultaneously with their associated trial events, but could have non-zero latencies. For example, pupil responses could anticipate predictable trial events. (3) Parameter recovery confirmed the validity of the fitting procedures, and we quantified the reliability of the parameter estimates for our sample dataset. (4) A cognitive task manipulation modulated pupil response amplitude. We provide our pupil analysis pipeline as open-source software (Pupil Response Estimation Toolbox: PRET) to facilitate the estimation of pupil responses and the evaluation of the estimates in other datasets.

## Introduction

Pupil size depends strongly on light levels, but it also covaries with an array of perceptual and cognitive processes – from attention to memory to decision making (for recent reviews, Binda & Gamlin, 2017; Ebitz & Moore, 2018; Mathôt, 2018; Wang & Munoz, 2015). Pupil size can be measured noninvasively and continuously, making pupillometry a promising tool for probing the ongoing dynamics linked to these processes. The pupil dilates in response to task-relevant stimuli (Hoeks & Levelt, 1993; Kang, Huffer, & Wheatley, 2014; Wierda, van Rijn, Taatgen, & Martens, 2012; Willems, Damsma, Wierda, Taatgen, & Martens, 2015; Willems, Herdzin, & Martens, 2015; Zylberberg, Oliva, & Sigman, 2012) and arousing, interesting, or surprising stimuli (Allen et al., 2016; Hess & Polt, 1960; Kloosterman et al., 2015; Knapen et al., 2016; Libby, Lacey, & Lacey, 1973; Nassar et al., 2012; Preuschoff, ‘t Hart, & Einhäuser, 2011), as well as in concert with internally-driven cognitive events, like mental calculation (Hess & Polt, 1964; Kahneman, Beatty, & Pollack, 1967), memorization (Kahneman & Beatty, 1966; Kang et al., 2014), and decision formation (Cheadle et al., 2014; de Gee et al., 2017; de Gee, Knapen, & Donner, 2014; Lempert, Chen, & Fleming, 2015; Murphy, Boonstra, & Nieuwenhuis, 2016; Murphy, Vandekerckhove, & Nieuwenhuis, 2014; Urai, Braun, & Donner, 2017; van Kempen et al., 2019). Pupil dilation is mediated by activity in the locus coeruleus (LC), hypothalamus, and superior colliculus, which interact with the pathways that control pupillary dilation and constriction (Mathôt, 2018; Wang & Munoz, 2015). The pupil time series may therefore carry information about multiple events within an experimental trial, as well as about anticipatory neural responses not available from behavioral reports alone, which provide retrospective rather than online measures.

A critical challenge in relating pupillary changes to specific perceptual and cognitive processes is that pupillary dynamics are relatively slow. Whereas perception and cognition unfold over timescales of a few hundred milliseconds, the pupil takes about 2 s to dilate and return to baseline in response to a single, brief stimulus (Hoeks & Levelt, 1993). However, the limiting factor in the speed of pupil dilation does not appear to be the dynamics of the neural responses that drive the pupil. For example, LC activity is tightly linked to pupil dilation (Aston-Jones & Cohen, 2005; de Gee et al., 2017; Joshi, Li, Kalwani, & Gold, 2016; Murphy, O’Connell, O’Sullivan, Robertson, & Balsters, 2014; Reimer et al., 2016; Varazzani, San-Galli, Gilardeau, & Bouret, 2015) and has much faster dynamics. LC neurons fire with a latency of ≤100 ms after a task-relevant stimulus, with a brief, phasic response (Aston-Jones & Cohen, 2005; Foote, Aston-Jones, & Bloom, 1980; Sara & Bouret, 2012). Therefore, the pupil size at a given time may reflect the influence of multiple preceding or ongoing internal signals related to distinct perceptual and cognitive processes. The standard approach to pupillometry, namely measuring the pupil size time series, cannot disentangle the influences of these various signals on the pupil size.

To address this challenge, researchers have begun to use models to link changes in pupil size to the distinct internal signals elicited by specific trial events (de Gee et al., 2017; de Gee et al., 2014; Hoeks & Levelt, 1993; Johansson & Balkenius, 2017; Kang et al., 2014; Kang & Wheatley, 2015; Knapen et al., 2016; Korn & Bach, 2016; Korn et al., 2017; Lempert et al., 2015; Murphy et al., 2016; Urai et al., 2017; van den Brink, Murphy, & Nieuwenhuis, 2016; van Kempen et al., 2019; Wierda et al., 2012; Willems, Damsma, et al., 2015; Willems, Herdzin, et al., 2015; Zylberberg et al., 2012). These signals can be thought of as the internal responses to trial events that drive pupil dilation, and the goal of modeling is to infer the properties of these internal signals, such as their amplitudes, from the pupil time series. Under constant luminance conditions, it is typical to model pupil dilations only, which are considered to be linked to internal signals that drive the sympathetic pupillary pathway (Mathôt, 2018).

Pupil response models typically incorporate two principles based on the work of Hoeks and Levelt (1993). First, the models assume a stereotyped pupil response function (puRF), which describes the time series of pupil dilation in response to a brief event. These authors found that the puRF is well described by a gamma function, and they reported average parameters for that function, which have been used in many studies (de Gee et al., 2017; de Gee et al., 2014; Kang et al., 2014; Kang & Wheatley, 2015; Lempert et al., 2015; Murphy et al., 2016; van Kempen et al., 2019; Wierda et al., 2012; Willems, Damsma, et al., 2015; Willems, Herdzin, et al., 2015; Zylberberg et al., 2012) – we refer to this specific form of the puRF as the “canonical puRF”. Second, the models assume that pupil responses to different trial events sum linearly to generate the pupil size time series; that is, they are general linear models (GLMs). This assumption is based on Hoeks and Levelt’s (1993) finding that, for the tested stimulus parameters, the pupil responded like a linear system. Incorporating these two principles, the pupil response to sequential trial events has been modeled as the sum of component pupil responses, where each component response is the internal signal time series associated with a single trial event convolved with the puRF. Using this approach, the pupil has been found to track decision periods (de Gee et al., 2017; de Gee et al., 2014; Lempert et al., 2015; Murphy et al., 2016; Murphy, Vandekerckhove, et al., 2014; van Kempen et al., 2019) and fluctuations in target identification during a rapid stimulus sequence (Wierda et al., 2012; Zylberberg et al., 2012) – findings that reveal the faster internal dynamics underlying the measured pupil time series.

Despite the promise of using such models to link distinct pupillary responses to specific trial events, there is currently no standard procedure for modeling the pupil time series. Hoeks and Levelt (1993) estimated the number, timing, and amplitudes of impulse signals that drove pupil dilations. Later studies assumed that every stimulus presentation was associated with a concurrent impulse signal and estimated only their amplitudes. Some studies have also included longer, cognitive events, like a decision period (de Gee et al., 2017; de Gee et al., 2014; Lempert et al., 2015; Murphy et al., 2016; van Kempen et al., 2019), or extra parameters to account for slow drifts in pupil size across the trial (Kang et al., 2014; Kang & Wheatley, 2015; Wierda et al., 2012; Willems, Damsma, et al., 2015; Willems, Herdzin, et al., 2015; Zylberberg et al., 2012). Most studies have used the canonical puRF (de Gee et al., 2017; de Gee et al., 2014; Kang et al., 2014; Kang & Wheatley, 2015; Lempert et al., 2015; Murphy et al., 2016; van Kempen et al., 2019; Wierda et al., 2012; Willems, Damsma, et al., 2015; Willems, Herdzin, et al., 2015; Zylberberg et al., 2012), but others used more complicated puRFs that could also capture pupil dilations and constrictions in response to changes in illumination (Korn & Bach, 2016; Korn et al., 2017), or that separately modeled transient and sustained components of the pupil dilation (Spitschan et al., 2017). Importantly, these pupil-modeling methods have not been systematically evaluated or compared, hindering the adoption of a field-wide standard.

Here we evaluate GLM procedures for modeling the pupil time series for trials with multiple rapid sequential events, under constant illumination. We conduct factorial model comparison to determine which parameters should be included, and we perform several validation and reliability tests using the best model. Based on the results, we recommend a specific model structure and fitting procedure for more general adoption and future testing. Critically, we found that timing parameters not typically included in pupil GLMs substantially improve model fits. Our findings indicate that puRF timing should be estimated for each observer, rather than assuming the canonical puRF, and that the internal signals driving the pupil response are not necessarily concurrent with stimulus onsets. We provide an open-source MATLAB toolbox, the Pupil Response Estimation Toolbox (PRET), which fits pupil GLMs to obtain event-related amplitudes and latencies, estimates parameter reliabilities, and compares models.

As a case study, here we analyzed data for a study on temporal attention – the prioritization of sensory information at specific points in time (Denison, Heeger, & Carrasco, 2017). Combining information about the expected timing of sensory events with ongoing task goals improves our perception and behavior (review by Nobre & van Ede, 2018). By studying the effects of temporal attention on perception, we can better understand the dynamics of visual perception. To understand these dynamics, a critical distinction must be made between temporal attention— prioritization of task-relevant time points—and temporal expectation—prediction of stimulus timing regardless of task relevance. Here, we manipulated temporal attention while equating expectation by using precues to direct voluntary temporal attention to specific stimuli in predictably timed sequences of brief visual targets (Denison et al., 2017; Denison, Yuval-Greenberg, & Carrasco, 2019; Fernandez, Denison, & Carrasco, 2019). Within the temporal attention dataset, we also compared two kinds of tasks—orientation discrimination and orientation estimation—which involved identical stimulus sequences and only differed in the required report. It is likely that estimation has a higher cognitive demand than discrimination, as it requires a precise response, as opposed to a two-alternative forced choice. Thus the physical stimuli were fixed while the cognitive demand varied between tasks. This dataset provided a good case study to evaluate GLM procedures for modeling the pupil time series as it had multiple rapid sequential events, required temporally precise cognitive control to attend to a relevant time point that varied from trial to trial, and included an orthogonal task manipulation that involved different cognitive demands for identical stimuli.

## Methods

### Data set

We reanalyzed eye-tracking data collected in a recent study on temporal attention by Denison, Heeger and Carrasco (2017). Thus behavioral procedures were identical to those previously reported (Denison et al., 2017; Denison et al., 2019). To maximize power of the pupil analysis, we combined the data from the two experiments in that study with identical stimulus sequences (Experiments 1 and 3). Experiment 1 used an orientation discrimination task, so we refer to it here as the Discrimination experiment. Experiment 3 used an orientation estimation task, so we refer to it here as the Estimation experiment. The stimuli were similar across experiments: on each trial, human observers were presented with a predictably timed sequence of two target gratings–which we refer to as T1 and T2–and judged the orientation of one of these gratings. An auditory precue before each sequence directed temporal attention to one or both grating times, and an auditory response cue after each sequence instructed observers which grating’s orientation to report.

### Observers

The observers were the same as in Denison et al. (2017), except that eye-tracking data from four observers in the Discrimination experiment could not be used for pupil analysis for technical reasons (e.g., excessive blinking). To better equate the number of observers in each experiment for the present study, we collected data from three new observers for the Discrimination experiment (as also reported in Denison et al., 2019). This gave 21 total observer data sets: 9 in Discrimination and 12 in Estimation. Three observers participated in both experiments, including author R.N.D. Therefore, 18 unique observers (10 female, 8 male; aged 19-43 years) are included in the present study. All observers provided informed consent, and the University Committee on Activities involving Human Subjects at New York University approved the experimental protocols. All observers had normal or corrected-to-normal vision.

### Stimuli

Stimuli were generated on an Apple iMac using Matlab and Psychophysics Toolbox (Brainard, 1997; Kleiner, Brainard, & Pelli, 2007; Pelli, 1997). They were displayed on a gamma-corrected Sony Trinitron G520 CRT monitor with a refresh rate of 100 Hz at a viewing distance of 56 cm. Observers’ heads were stabilized by a head rest. A central white fixation “x” subtended 0.5° visual angle. Visual target stimuli were 4 cpd sinusoidal gratings with a 2D Gaussian spatial envelope (standard deviation 0.7°), presented in the lower right quadrant of the display centered at 5.7° eccentricity (**Figure 1a**). Stimuli were high contrast (64% or 100%, which we combined as there were no behavioral differences). Placeholders, corners of a 4.25° × 4.25° white square outline (line width 0.08°) centered on the target location, were present throughout the display to minimize spatial uncertainty. The stimuli were presented on a medium gray background (57 cd/m2). Auditory precues were high (precue T1: 784 Hz; G5) or low (precue T2: 523 Hz; C5) pure sine wave tones, or their combination (neutral precue). Auditory stimuli were presented on the computer speakers.

**Figure 1.**
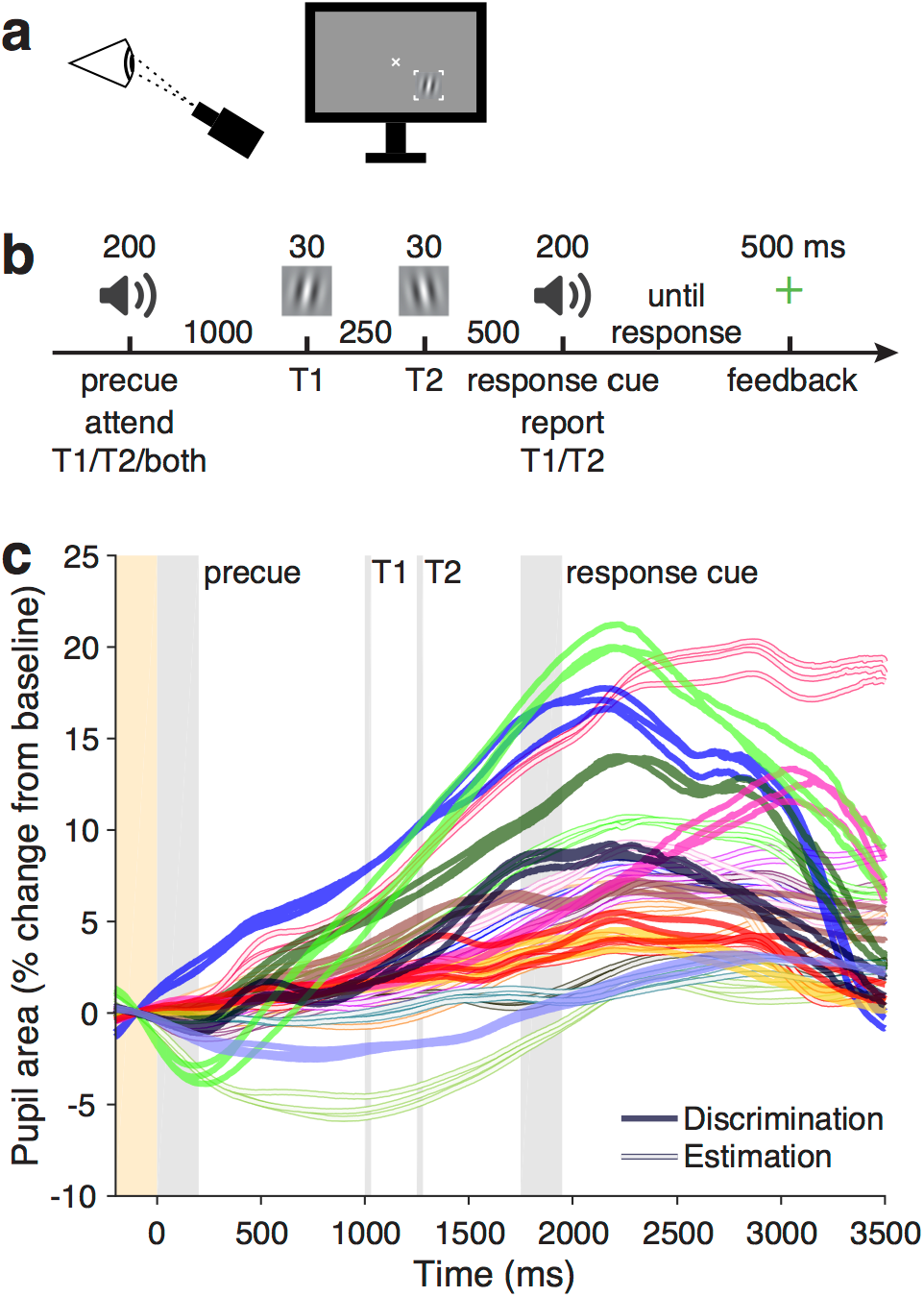
Task and pupil time series. **a)** Setup of visual display and eye tracker. **b)** Trial sequence. In the Discrimination task, observers reported whether the probed stimulus was tilted CW or CCW. In the Estimation task, a response grating (not shown) appeared after the response cue, and observers adjusted it to report the exact orientation of the probed stimulus. **c)** Pupil time series (colored lines), mean across trials in each condition for each observer. Filled lines are observers in the Discrimination experiment, and open lines are observers in the Estimation experiment. Each observer has a unique color, and three observers participated in both experiments (same color filled and empty). Three lines per observer and experiment show different precuing conditions (precue T1, T2, neutral), i.e., independent sets of trials. Time series were baseline-normalized per trial (baseline period shaded yellow). Gray shaded regions are trial events.

### Behavioral procedures

#### Basic task and trial sequence

Observers judged the orientation of grating patches that appeared in short sequences of two target stimuli per trial (T1 and T2). Targets were presented for 30 ms each at the same spatial location, separated by stimulus onset asynchronies (SOAs) of 250 ms (**Figure 1b**). An auditory precue 1000 ms before the first target instructed observers to attend to one or both of the targets. Thus there were three precue types: attend to T1, attend to T2, or attend to both targets. Observers were asked to report the orientation of one of the targets, which was indicated by an auditory response cue 500 ms after the last target. The duration of the precue and response cue tones was 200 ms. The timing of auditory and visual events was the same on every trial.

#### Discrimination task

In the Discrimination experiment, observers performed an orientation discrimination task (**Figure 1b**). Each target was tilted slightly clockwise (CW) or counterclockwise (CCW) from either the vertical or horizontal axis, with independent tilts and axes for each target, and observers pressed a key to report the tilt (CW or CCW) of the target indicated by the response cue, with unlimited time to respond. Tilt magnitudes were determined separately for each observer by a thresholding procedure before the main experiment. Observers received feedback at fixation (correct: green “+”; incorrect: red “-”) after each trial, as well as feedback about performance accuracy (percent correct) following each experimental block.

#### Estimation task

In the Estimation experiment, observers performed an orientation estimation task (**Figure 1b**). Target orientations were selected randomly and uniformly from 0-180°, with independent orientations for each target. Observers estimated the orientation of the target indicated by the response cue by adjusting a grating probe to match the perceived target orientation. The probe was identical to the target but appeared in a new random orientation. Observers moved the mouse horizontally to adjust the orientation of the probe and clicked the mouse to submit the response, with unlimited time to respond. The absolute difference between the reported and presented target orientation was the error for that trial. Observers received feedback at fixation after each trial (error <5°, green “+”; 5-10°, yellow “+”; ≥10°, red “-”). Additional feedback after each block showed the percent of trials with <5° errors, which were defined to observers as “correct”.

#### Training and testing sessions

All observers completed one session of training prior to the experiment to familiarize them with the task and, in the Discrimination experiment, determine their tilt thresholds. Thresholds were selected to achieve ∼79% performance on neutral trials. Observers completed 640 trials across 2 one-hour sessions. All experimental conditions were randomly interleaved across trials.

### Eye data collection

Pupil size was continuously recorded during the task at a sampling frequency of 1000 Hz using an EyeLink 1000 eye tracker (SR Research). Raw gaze positions were converted into degrees of visual angle using the 5-point-grid calibration, which was performed at the start of each experimental run. Online streaming of gaze positions was used to ensure central fixation (<1.5° from the fixation cross center) throughout the experiment. Initiation of each trial was contingent on fixation, with a 750 ms minimum inter-trial interval. Observers were required to maintain fixation, without blinking, from the onset of the precue until 120 ms before the onset of the response cue. If observers broke fixation during this period, the trial was stopped and repeated at the end of the block.

### Preprocessing

Data files from the eye tracker were imported to Matlab to perform all preprocessing and modeling with custom software. The raw time series from each session was epoched into trials spanning from −500 to 3500 ms, relative to the precue at 0 ms. Blinks were interpolated trial by trial using a cubic spline interpolation method (Mathôt, 2013). All trials were individually baseline normalized by calculating the average pupil size over the region from −200 to 0 ms, then calculating the percent difference from this baseline at each point along the time series:

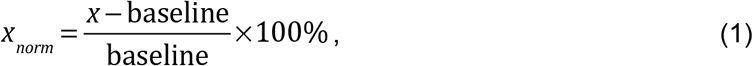

where *x*_*norm*_ is the normalized data and *x* is the raw data. We normalized the time series in this way to obtain meaningful units of percent change from baseline, but we note there are also arguments for a purely subtractive baseline correction procedure (Mathôt, Fabius, Heusden, & Stigchel, 2018; Reilly, Kelly, Kim, Jett, & Zuckerman, 2018). Trials were grouped into conditions depending on the precue (T1, T2, neutral), and the mean time series was calculated across trials in each condition for each observer.

### Pupil modeling

#### GLM modeling framework

Measured pupil size time series were modeled as a linear combination of component pupil responses (**Figure 2**). A component pupil response is the predicted pupil size time series associated with a single internal (neural) signal that leads to pupil dilation. Mathematically, an internal signal was represented as a time series concurrent with the measured pupil size time series. The component pupil response for a given internal signal was calculated by convolving the signal time series with a pupil response function. The general pupil response function takes the form

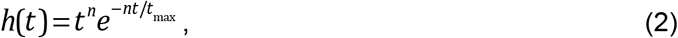

where *h* is the pupil size, *t* is the time in ms, *n* controls the shape of the function, and *t*_max_ controls the temporal scale of the function and is the time of the maximum (Hoeks and Levelt 1993) (**Figure 2a**). For a given measured pupil size time series, each internal signal was convolved with the same pupil response function. Each component pupil response was assumed to be dilatory.

**Figure 2.**
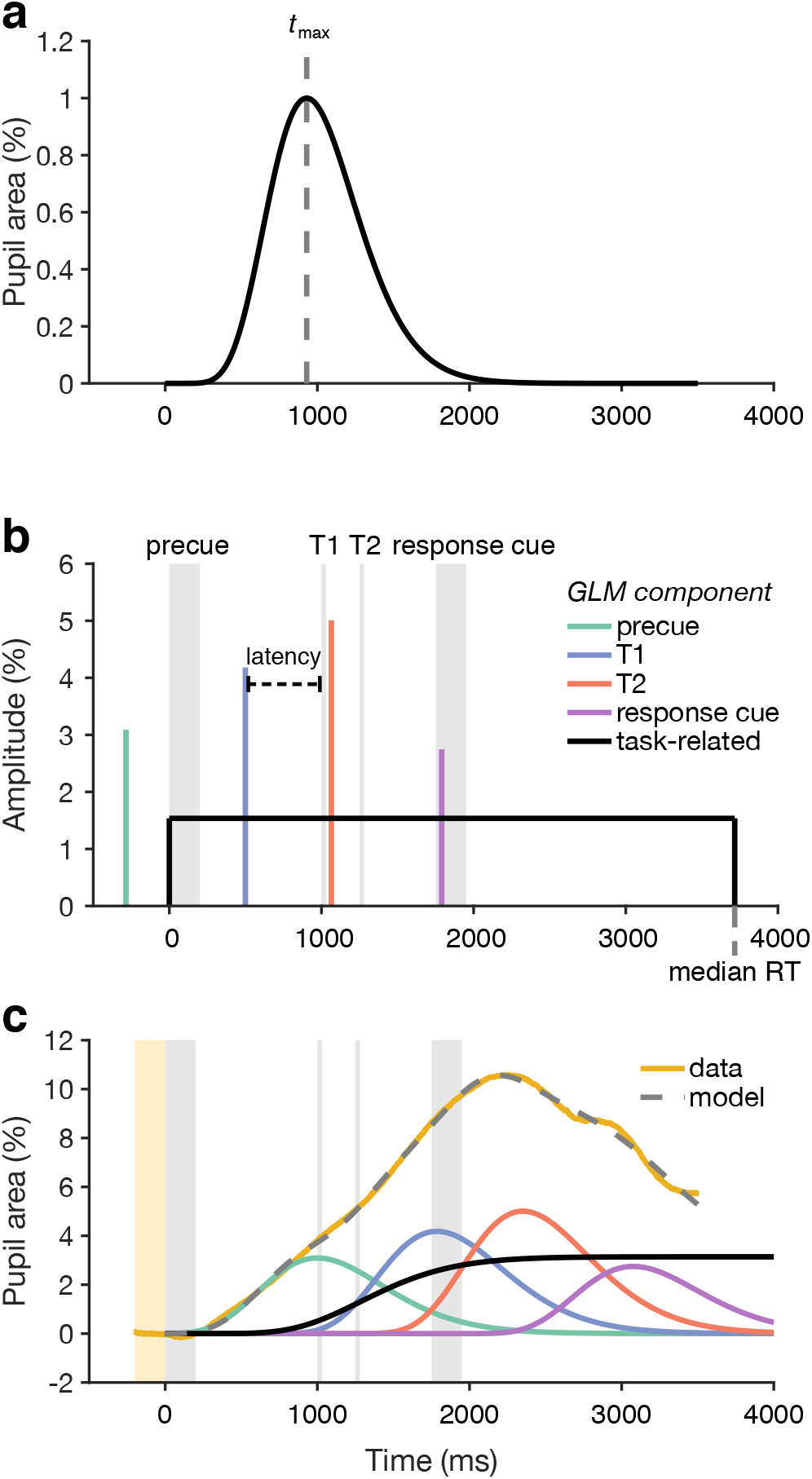
General linear modeling of the pupil time series. **a)** Pupil response function (puRF), which describes the pupillary response to an impulse event. The canonical puRF, an Erlang gamma function with *n* = 10.1 and *t*_*max*_ = 930 ms (vertical dashed line), is shown. **b)** Internal signals hypothesized to drive pupil dilation. The internal signal associated with each trial event (brief auditory and visual stimuli, gray shaded regions) is modeled as a delta function (vertical colored lines) with some amplitude and some latency with respect to the event. A sustained, task-related signal could also be modeled. Shown here is a boxcar (black line), which starts at the onset of the precue and lasts until the median RT of the modeled trials. **c)** The mean pupil time series across trials (yellow line) is modeled in two steps. First, each internal signal time series is convolved with the puRF to form component pupil responses (colored lines, legend in panel b). Second, the component pupil responses are summed to calculate the model prediction (gray dashed line). Parameters of the model, such as the amplitudes and latencies of the internal event signals, are fit using an optimization procedure.

We assumed there was a transient internal signal (de Gee et al., 2014; Hoeks & Levelt, 1993; Wierda et al., 2012) associated with each event in the trial sequence: the precue, T1, T2, and the response cue. Each of these event-related signals took the form of a Dirac delta function. An additional internal signal could be included to model a constant, sustained signal associated with task engagement (de Gee et al. 2014). This task-related signal took the form of a boxcar function, with nonzero values starting at the precue and ending at the median response time of the observer being modeled. Thus, we modeled our measured pupil size time series as the linear combination of up to four event-related and one task-related component pupil responses.

#### Model parameters

We fit models of up to eleven parameters to a given pupil size time series. The possible parameters were: internal signal amplitudes and latencies for each trial event; internal signal amplitude for the task-related response or alternatively a slope parameter specifying a linear drift across the trial; one parameter specifying the timing of the pupil response function; and one baseline shift parameter.

Each event-related signal had an amplitude parameter and a latency parameter associated with it. The amplitude parameter was the value of the nonzero point of the delta function and indicated the magnitude of the internal signal, and thus determined the magnitude of the component pupil response associated with it. The latency parameter was the time (in ms) of the nonzero value, relative to the time of its corresponding event. The task-related signal only had an amplitude parameter associated with it because it was assumed to start at the beginning of the trial.

The pupil response function that was convolved with each signal had two parameters: *t*_max_, which controls the temporal scale and time of the peak, and *n*, which controls the shape of the function. Only *t*_max_ was estimated while *n* was set to the canonical value of 10.1 (Hoeks & Levelt, 1993). The *t*_max_ parameter can be interpreted as the time it takes an observer’s pupil to dilate maximally in response to an internal signal. The pupil response function was normalized such that the event-related and task-related amplitude parameters indicated the percentage increase in pupil size attributable to the corresponding signal. The pupil response function was normalized to a maximum value of 1, so that an amplitude value of 1 corresponded to a 1% increase in pupil size from baseline. For the task-related amplitude, the puRF was normalized such that the puRF convolved with the boxcar had a maximum value of 1. Thus, a task-related amplitude of 1 also corresponded to a 1% increase in pupil size from baseline to peak size.

The final parameter was a baseline shift parameter we termed the *y*-intercept (*y*-int). The *y*-int parameter was simply a shift along the y-axis of the entire predicted pupil size time series. We included this in the model because we noticed that for some observers, although all trials were baseline-normalized during preprocessing based on a time window before the precue, pupil size was decreasing during this window and continued to decrease until shortly after the precue. This meant that pupil dilations during the trial sequence started from a value below the calculated baseline. Without accounting for this shift in baseline with the *y*-int parameter, the model would underestimate the amplitude of the component pupil responses.

#### Model comparison and selection

Previous linear models based on the pupil response function only estimated the amplitude of component pupil responses (e.g., de Gee et al., 2014; Wierda et al., 2012). The component responses were assumed to onset at the time of the corresponding trial event, and pupil response functions were assumed to be identical across observers. However, these assumptions have never been systematically evaluated. Here, we asked whether introducing additional timing parameters would allow more accurate modeling of the pupil response. In addition, the characteristic slow pupil dilation throughout a trial has been modeled in different ways, as either a linear drift (Wierda et al., 2012) or a sustained task-related response convolved with the pupil response function (de Gee et al., 2014). We compared these two possibilities.

We compared 24 different models (**Table 1**). These models included all permutations of latency, *t*_max_, and *y*-int as fixed vs. free parameters and three different forms of the task-related component. Fixed values are reported in **Table 1**. The three task-related components tested were a boxcar function convolved with the pupil response function, a linear function, and no task-related component. The boxcar function was nonzero from 0 ms (the onset of the precue) to the median response time for a given observer and had one amplitude parameter for the height of the boxcar. The linear function had a slope parameter and a fixed intercept of 0%. It represented a linear drift throughout the whole trial and did not depend on response time. Amplitude parameters for each trial event were always estimated. Each model was fit to the mean time series for each condition and observer. We also checked that the results held when fitting to the single trial time series; in this case, baseline-corrected single trial time series were concatenated and fit to concatenated model time series.

**Table 1.**
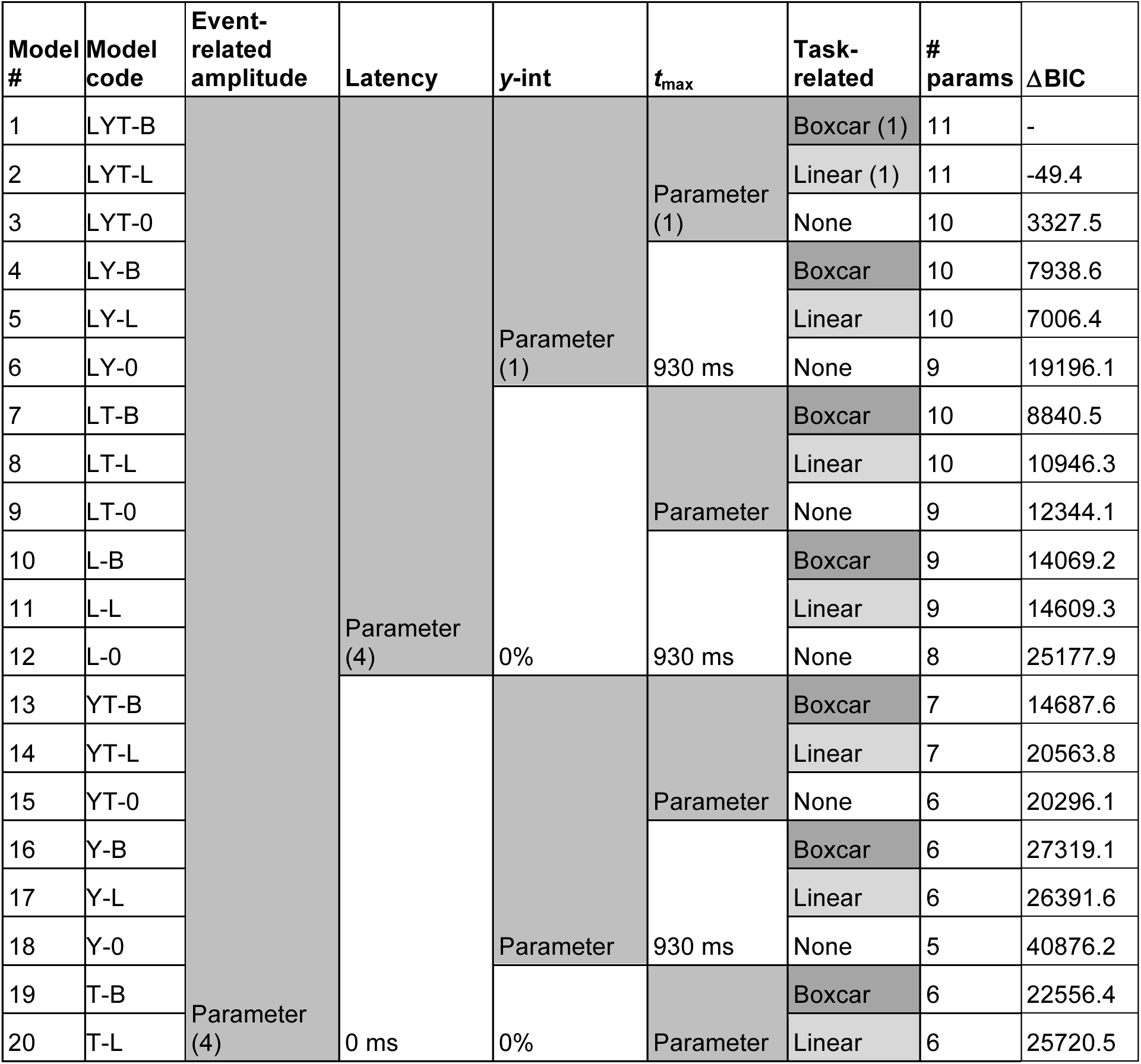
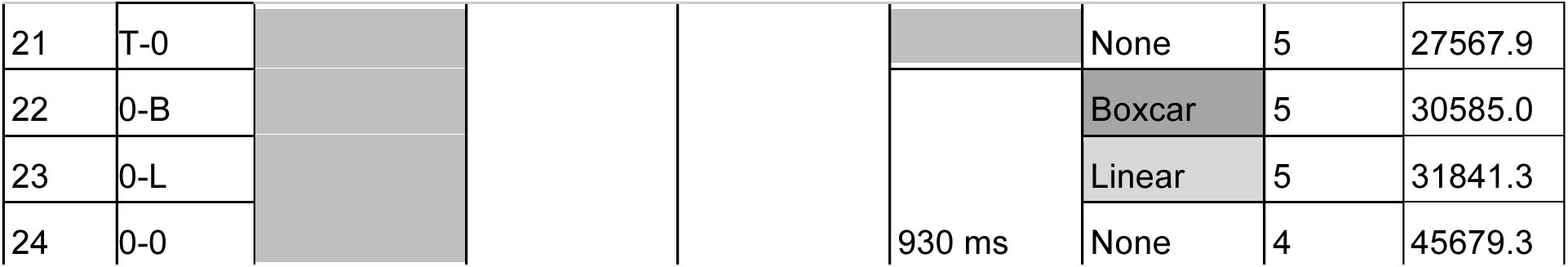
Factorial model comparison. All models tested are shown with their parameters. Rows are models; columns including gray shading are factors. Gray shading indicates parameters were fit, with number of parameters in parentheses. No shading indicates parameters were fixed, with the fixed value given. Model code: L = latency, Y = *y*-int, T = *t*_max_, 0 = none; task component: B = boxcar, L = linear, 0 = none.

To compare models, the Bayesian Information Criterion (BIC) was calculated for each model and observer across conditions and averaged at the group level to get one metric per model. The model with the lowest metric for most observers was selected as the best model and used for further analysis. The BIC was chosen as the comparison metric to account for the differing numbers of parameters among models.

We also assessed cross-validated *R*^2^ for each model using a 4-fold cross-validation procedure, in which models were fit to 75% of the data and tested on the remaining 25%. Cross-validated *R*^2^ was calculated by comparing the model prediction to the mean across trials of the held-out data for each fold and averaging across folds. A noise ceiling for *R*^2^ was calculated by comparing the mean across trials of the fitted data to the mean across trials of the held-out data for each fold and averaging across folds. *R*^2^ values were computed for each model for each observer and then averaged across observers.

#### Parameter estimation

Model parameters were estimated for the mean pupil time series for each condition and observer using a two-phase procedure. In both phases, the cost function for determining goodness of fit was defined as the sum of the squared errors between the measured pupil time series and the predicted (model-generated) pupil time series in the time window from 0 to 3500 ms. First, the cost function was evaluated at 2000 sets of parameter values sampled from independent uniform distributions of each parameter. These distributions were bounded by the parameter constraints described below. Second, constrained optimizations (MATLAB *fmincon*) were performed starting from the 40 sets of parameter values with the lowest cost from the first phase. The set of optimized parameter values minimizing the cost function was selected as the best estimate.

Parameter constraints were selected to ensure parameters could vary meaningfully within physiologically feasible ranges. Event and task-related amplitudes were constrained to the range of 0 to 100, meaning that any single internal signal could not evoke a pupil response in excess of 100 percent change from baseline. This range also enforces that pupil responses refer to dilation as opposed to constriction. Event-related latencies were constrained to a range of −500 to 500 ms. The *t*_max_ value was constrained to a range of 0 to 2000 ms. The *y*-int parameter was constrained to a range of −20 to 20 percent change from baseline. The slope parameter during model comparison was constrained to a range of 0 to 50 percent change over the trial window of 3500 ms.

#### Bootstrapping procedure for parameter estimation

To obtain robust parameter estimates, and to quantify the uncertainty in the parameter estimates due to both noise in the data and variability in the fitting procedure, we used bootstrapping. Parameter estimation was performed on 100 bootstrapped mean time series for each condition and observer. Each bootstrapped time series was formed by randomly resampling the underlying set of trials with replacement, with the number of trials per sample equal to the original number of trials. This produced 100-element distributions of each model parameter for each condition and observer. The median of a given parameter distribution was taken as the bootstrapped estimate of that parameter. The uncertainty of this estimate was quantified as the 95 percent confidence interval.

#### Parameter recovery

To evaluate the parameter estimation procedure, we fit the model to artificial data generated by the model, with no noise. Because the form of the noise in pupil data is unknown, we relied on the bootstrapping procedure (in which the noise comes from the data itself) to quantify the uncertainty of parameter estimates and performed parameter recovery only to verify the accuracy of the fitting procedure and check for redundancies within the model structure. A set of 100 artificial time series was simulated by generating 100 sets of model parameters independently sampled from uniform distributions and calculating the resulting time series for each. Event and task-related amplitude values were sampled from a range of 0 to 10%, latency values were sampled from −500 to 500 ms, *t*_max_ values were sampled from 500 to 1500 ms, and *y*-int values were sampled from −4 to 4%. Response time values used in the boxcar function for the task-related component varied from 2350 to 3350 ms. The parameter estimation procedure was performed on each artificial time series (without bootstrapping), producing an output set of parameters for each input set of parameters. To evaluate the parameter recovery, input parameters were plotted against output parameters and the Pearson correlation coefficient was calculated.

### Statistical testing

Hypothesis testing was performed using the median of the parameter estimates from the bootstrapping procedure. A linear mixed effects model was used to analyze the combined data across two experiments, each with a within-observer design, and in which three observers completed both experiments. A linear mixed effects model was created using the *lme4* package in R, with experiment and precue condition as fixed effects and observer as a random effect. We tested for main effects and interactions by approximating likelihood ratio tests to compare models with and without the effect of interest.

## Results

We evaluated the ability of general linear models (GLMs) to capture pupil area time series during experimental trials with rapid sequences of events. We tested the model on a sample data set in which four sequential stimuli were presented within 2.25 s on each trial (**Figure 1a,b**, see Methods). Given that 2 s is the approximate length of a typical pupil impulse response function (Hoeks & Levelt, 1993), we asked whether pupil responses to the successive events within a trial can be meaningfully recovered and what model form would best describe the pupil area time series over the course of a trial.

The data set included two experiments with the same stimulus sequence but different types of behavioral reports (orientation discrimination or orientation estimation) at the end of each trial. The experiments were previously reported with only the behavioral analysis (Denison et al., 2017) and with microsaccade analysis (Denison et al., 2019). Pupil area time series, averaged across trials for each experimental condition, were highly consistent within individual observers but varied considerably across observers (**Figure 1c**), motivating an individual-level modeling approach.

The modeling framework assumed that the pupil response time series is a linear combination of component responses to various trial events, along with a task-related pupil response in each trial (**Figure 2**) (de Gee et al., 2014; Hoeks & Levelt, 1993; Wierda et al., 2012). Each trial event was modeled as an impulse of variable amplitude (**Figure 2b**), which was convolved with a pupil response function (**Figure 2a**) to generate the corresponding component response (**Figure 2c**). The sum of all component responses was the predicted pupil time series (**Figure 2c**).

### Model comparison: Timing parameters improve fits

We compared 24 alternative models to determine what model structure would allow the best prediction of the pupil response time series. In particular, we asked whether the addition of two timing parameters would improve model fits over that of the standard model. The first timing parameter was event latency: a trial event impulse could have a non-zero latency with respect to its corresponding event, rather than being locked to the event onset. The second timing parameter was *t*_max_: the time-to-peak of the pupil impulse response function could vary across individuals. We also tested different forms of the sustained, task-related pupil response and the inclusion of a baseline parameter (*y*-int) to account for differences not removed by pre-trial baseline normalization. We used factorial model comparison (Keshvari, van den Berg, & Ma, 2012; Ma, 2018; van den Berg, Awh, & Ma, 2014) to test the contribution of each of these parameters to predicting pupil response time series (**Table 1**).

All four tested parameters (event latency, *t*_max_, task-related component, and *y*-int) significantly improved model fits (**Figure 3**; multi-way within-observers ANOVA on BIC scores, main effects of latency: F(1,20) = 143.39, p = 1.4e-10, mean across observers and models ΔBIC = − 5465.681; *t*_max_: F(1,20) = 76.59, p = 2.8e-08, ΔBIC = −10324.07; task-related: F(2,40) = 19.88, p = 1.0e-06, ΔBIC (box minus linear) = −1379.174, ΔBIC (box minus none) = −8558.598; *y*-int: F(1,20) = 20.53, p = 2.0 e-04, ΔBIC = −6865.328). We also observed interactions between some factors including task-related component by *t*_max_: F(2,40) = 23.66, p = 1.7e-07; tmax by *y*-int: F(1,20) = 8.27, p = 9.4e-03; task-related component by latency: F(2,40) = 3.25, p = 4.9e-02; *t*_max_ by latency: F(1,20) = 7.00, p = 1.6e-02. The best model for most observers (model 1) included all the tested parameters, with the task-related component modeled as a boxcar convolved with the pupil response function. The model fit the mean time series data well (mean across observers, *R*^2^ = 0.9871; cross-validated *R*^2^ = 0.6737, 98% of noise ceiling). The single-trial *R*^2^ was 0.1983; this value reflects the noise at the single-trial level. Model 2, in which the task-related component was modeled as a linear function of time but which was otherwise identical to model 1, performed similarly (*R*^2^ = 0.9895; cross-validated *R*^2^ = 0.6761, 98% of noise ceiling; single-trial *R*^2^ = 0.1990). These two types of task-related components also performed similarly at the factor level (ΔBIC = −1379.174). Otherwise, model 1 was significantly better than every other model (paired t-tests of model 1 BIC vs. each model’s BIC, all t > 3.09, all p < 5.78e-03 uncorrected; with Bonferroni correction for 23 pairwise comparisons, all but model 3 had p < 0.05). We found a similar pattern across models for cross-validated *R*^2^ as well as when we fit to single-trial data. Model 1 was consistently the best model, and latency, *t*_max_, and task-related parameters improved model fits. Therefore, the addition of timing parameters to the standard model substantially improved model fits.

**Figure 3.**
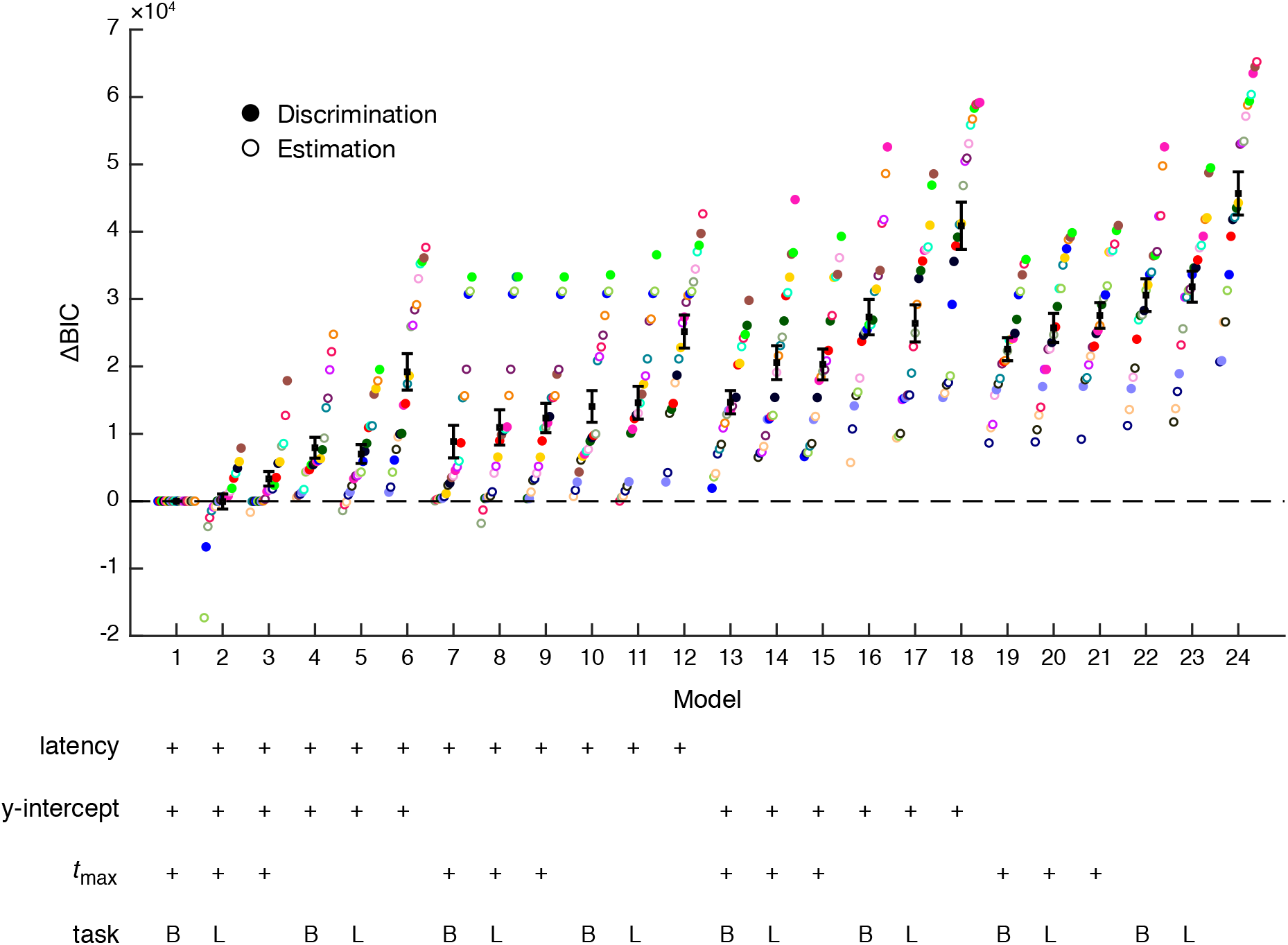
Model comparison. Difference in BIC score for each observer and model with respect to model 1, the best model for most observers. Each color corresponds to an individual observer. Black square with error bar corresponds to mean difference in BIC score across all observers. Crosses in the table indicate that the parameter type was fit for that model. In the task row, B indicates boxcar and L indicates linear function.

Our final test of the model’s structure was to ask whether modeling all trial events was needed to predict the pupil area time series. In particular, the two visual target events were separated by only 250 ms, which is short compared to the dynamics of the pupil response. To test whether the model captured separate pupil responses to the two targets, we compared models that included either two target events (model 1) or only one target event. The two-target model outperformed the one-target model, t(20) = 4.23, p = 4.1e-04, consistent with separate pupil responses to the two rapidly presented targets.

### Validation of the fitting procedure: Parameter recovery and tradeoffs

We evaluated the best model (model 1) and fitting procedures in several ways. First, we sought to validate the model and fitting procedures by performing parameter recovery on simulated data. Redundancies in the model or lack of precision to resolve the unique contributions of different trial events to the pupil time series would result in parameter tradeoffs, and noise in the fitting procedure from stochastically searching a high-dimensional parameter space would result in variability in the parameter estimates. We performed parameter recovery to assess these possibilities by generating 100 simulated time series with known parameter values and then fitting the model. Parameter estimates from simulated data tended to be similar to the true values for the entire range of tested values (strong diagonals on the 2D histograms in **Figure 4** and correlations in each panel). This was also the case when the range of T1-T2 SOAs was restricted to ±100 ms around the experimental SOA of 250 ms (**Figure S1**), as well as when noise on single trials was simulated (**Figure S2**). Parameter recovery accuracy was lowest for the T1 and T2 amplitudes, likely because of their close temporal proximity. Accuracy for the timing parameters was generally high.

**Figure 4.**
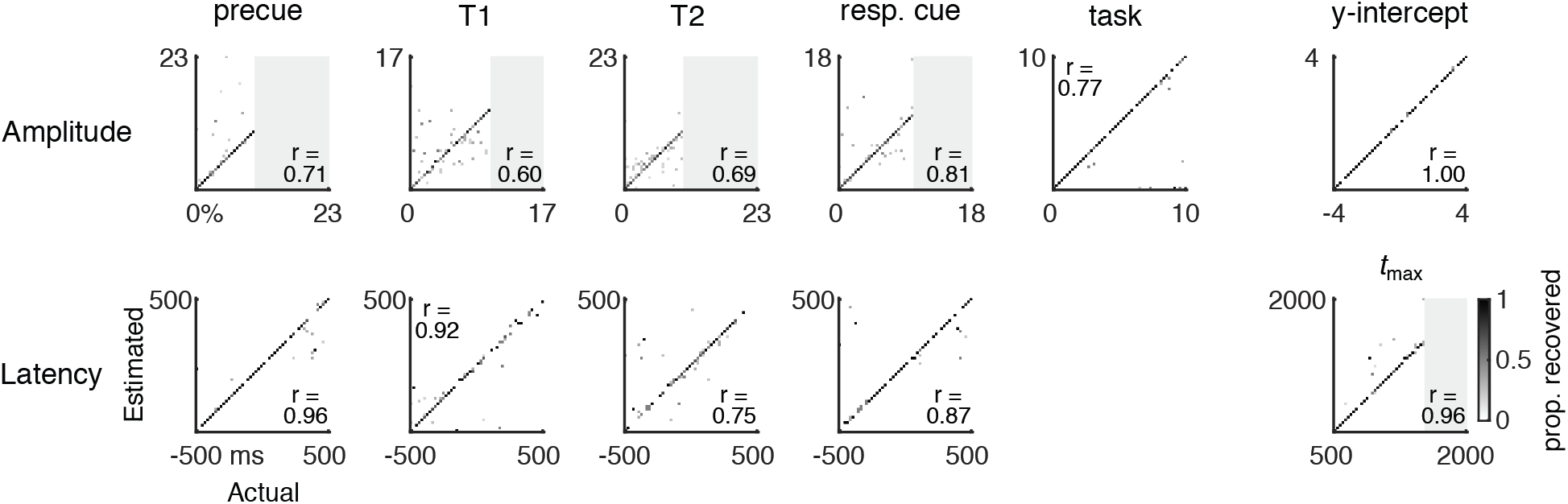
Parameter recovery for simulated data. 2D histograms showing the proportion of model fits for which a given actual parameter value (x-axis) was fit as a given estimated parameter value (y-axis). Perfect model recovery would appear as a diagonal black line (all actual values recovered as the same estimated values). Each panel is one parameter. Pearson correlations between actual and estimated values are given in each panel. Gray shaded regions are outside the range of actual parameter values simulated.

We assessed parameter tradeoffs first by examining correlations between estimated values for pairs of parameters in the simulated data. No correlations were significant after Bonferroni correction for multiple comparisons (**Figure S3**). The lack of significant correlations between parameters in the simulated data indicates that parameter tradeoffs are not inherent to the structure of the model or the fitting procedure.

We next assessed parameter tradeoffs by examining correlations between pairs of parameter values (bootstrap medians) estimated from the real data. All event-related amplitudes were positively correlated with each other (r > 0.61, p < 1.2e-07). Certain event latencies were also positively correlated with each other (T1 with T2, T2 with response cue; r > 0.44, p < 2.7e-04). These correlations are likely to arise from true statistical dependencies in the data; e.g., some observers have generally stronger pupil dilation responses, across all events. In addition to these positive correlations, T1 latency negatively correlated with event-related amplitudes (precue, T1, T2; r < −0.44, p < 3e-04) and precue latency negatively correlated with task-related amplitude (r = −0.48, p < 6.01e-05). *t*_max_ negatively correlated with *y*-int (r = −0.43, p = 4.73e-04) and positively correlated with response cue latency (r = 0.53, p = 6.4e-06). The negative correlations could arise from parameter tradeoffs driven by noise in the real data, or from true statistical dependencies in the pupil responses.

### Reliability of parameter estimates for individual observers

We next evaluated the reliability of the parameter estimates in real data. Real data have multiple sources of noise, some of which are unknown, so to estimate the reliability of parameter estimates given such noise, we used a bootstrapping procedure. This procedure allowed us to estimate the reliability of parameter estimates for individual observers. Parameter estimates and their reliabilities for an example observer are shown in **Figure 5**. We define “reliability” as the range of the 95% confidence interval (CI). The mean confidence interval for each parameter estimate is shown in **Figure 5**. The mean reliability of different event types were: trial event amplitude: 2.2118%; task-related amplitude: 2.7544%; trial event latency: 342.1034 ms; *t*_max_: 334.3857 ms; *y*-int: 0.74%. Due to this variability across bootstrap samples, we recommend using the median of the bootstrapped distribution as a robust parameter estimate, and we adopt this practice going forward.

**Figure 5.**
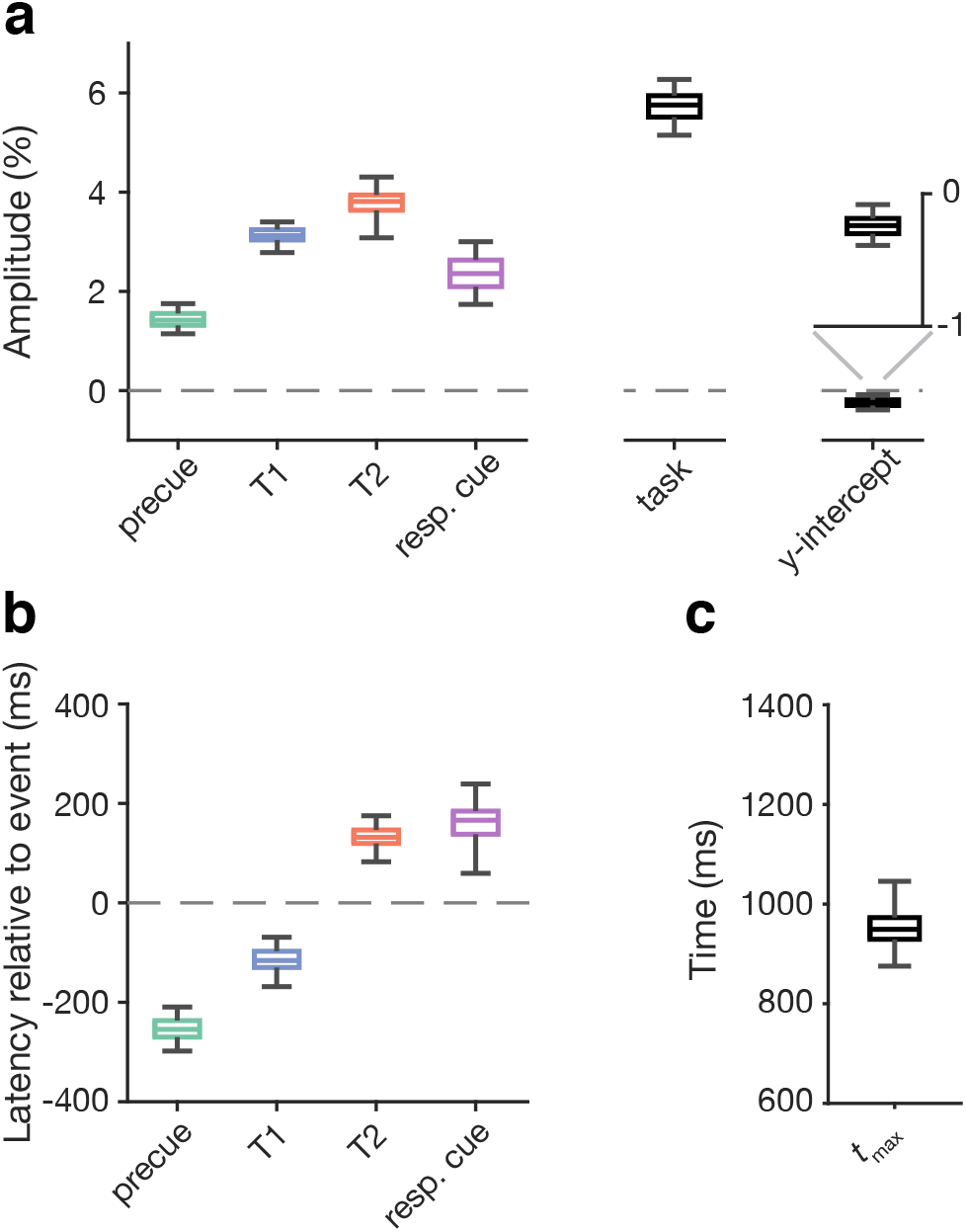
Reliability of individual parameter estimates for a representative observer. **a)** Amplitude and *y*-int estimates. Box and whisker plots show bootstrap median along with 50% and 95% bootstrapped confidence intervals. **b)** Latency and **c)** *t*_max_ estimates, plotted as in panel a.

### Consistency of parameter estimates within observers and variability across observers

We next investigated the consistency of parameter estimates within each observer as well as their variability across observers. To do this, we measured the consistency of parameter estimates for a given observer across independent sets of trials (**Figure 6**). We split the trials for each observer based on the experimental condition (3 conditions per observer: precue T1, T2, neutral). Parameter estimates were generally consistent for individual observers, though latency was less consistent than the other parameters. In contrast to the within-observer consistency, all parameters varied substantially across observers. In particular, the *t*_max_ parameter, which describes the dynamics of the pupil response function, was highly consistent within individual observers but varied over a large range across observers (from ∼700 to ∼1600 ms). These findings underscore the importance of modeling individual pupil response time series rather than only considering a group average time series, and they further show the importance of modeling individual observer pupil response dynamics rather than assuming a fixed pupil impulse response function.

**Figure 6.**
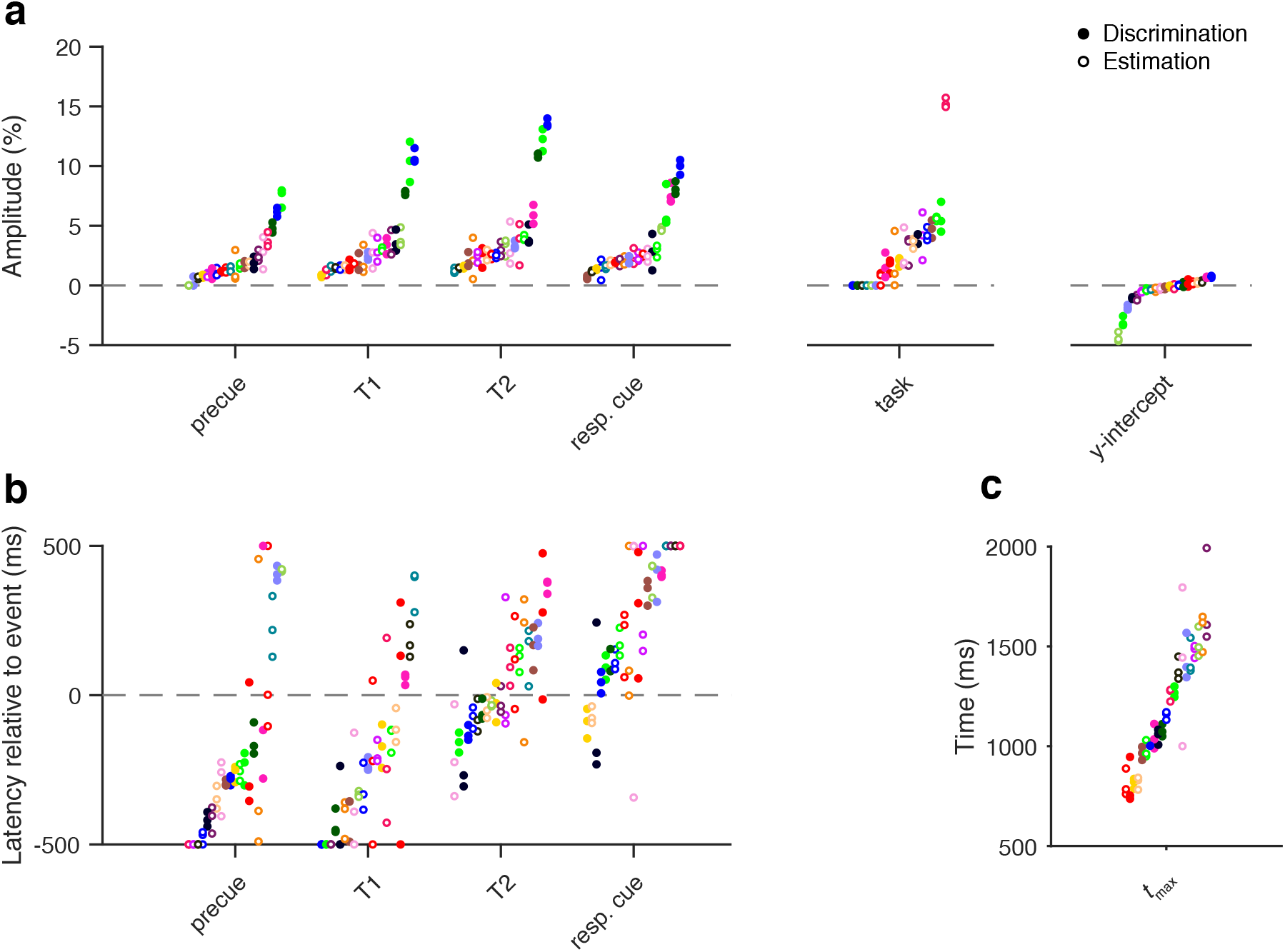
Consistency of parameter estimates across independent sets of trials. **a)** Amplitude and *y*-int estimates. Each point is one condition (precue T1, T2, neutral) for one observer; each condition was fit separately. Each observer has a unique color. Filled points are from the Discrimination experiment, empty points are from the Estimation experiment. **b)** Latency and **c)** *t*_max_ estimates, plotted as in panel a.

### Parameter estimates for the temporal attention experiment: Amplitude depends on task

To demonstrate how modeling can reveal cognitive modulations of the pupil response, we used the developed modeling procedure to evaluate the parameter estimates in the experimental data. We calculated separate estimates for each precue type (T1, T2, neutral), separately for the Discrimination and Estimation experiments, which had the same stimulus sequence but different types of behavioral reports (**Figure 7**). No differences were found among the different precue types (*χ*^2^(2) < 5.77, p > 0.05 for all parameters). There were also no interactions between precue and experiment (*χ*^2^(2) < 3.70, p > 0.15). So here we report the mean across precues for each experiment.

**Figure 7.**
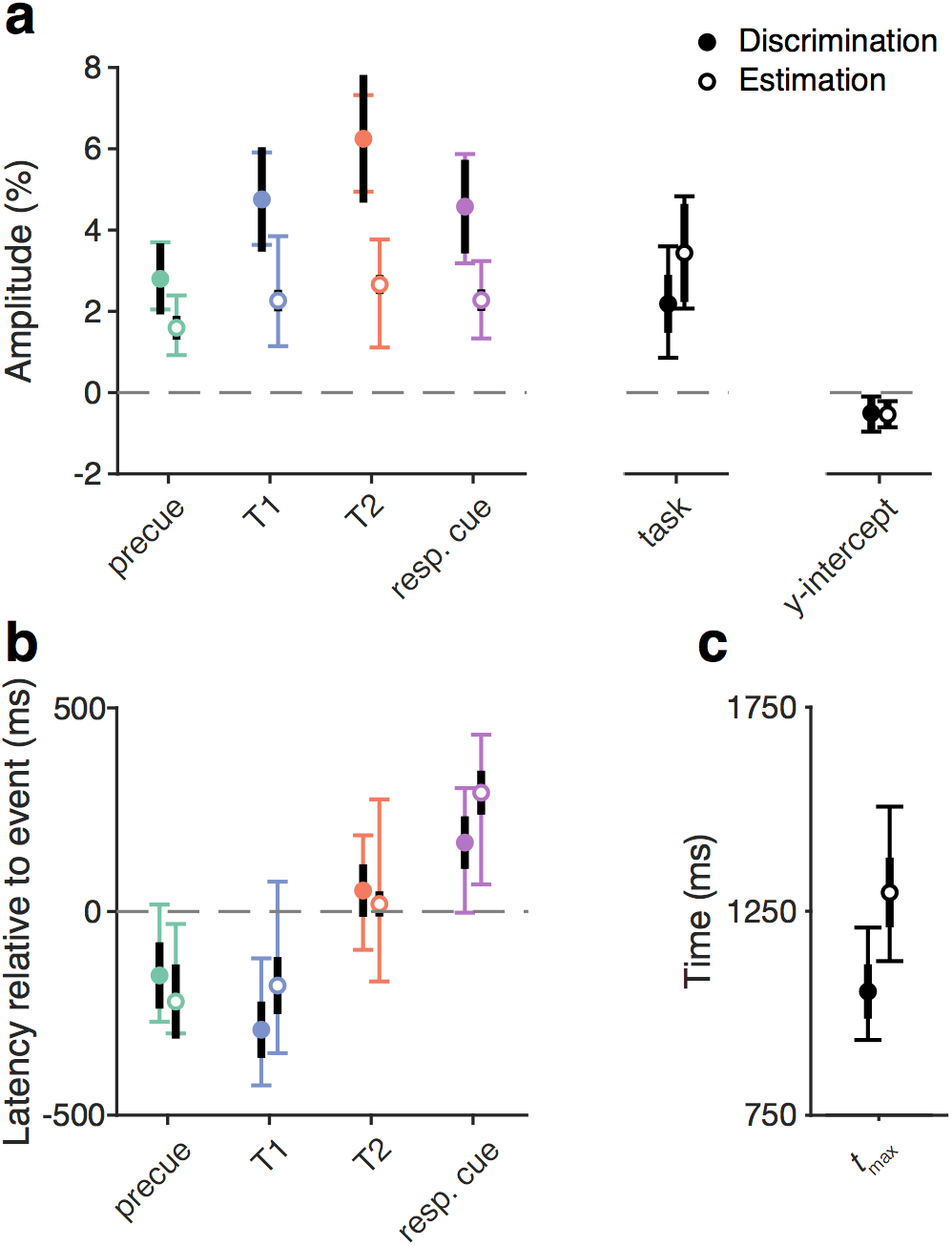
Group parameter estimates. **a)** Amplitude and *y*-int estimates. Colored points show mean of bootstrap medians across observers and conditions. Filled points are from the Discrimination experiment, empty points are from the Estimation experiment. Thin colored error bars show mean 95% confidence intervals across observers. Thick black error bars show SEM across observers. **b)** Latency and **c)** *t*_max_ estimates, plotted as in panel a.

Amplitude estimates for trial events were 1-7% change from baseline, and the amplitude of the decision-related signal was 2.1839% (Discrimination) or 3.4369% (Estimation). The mean *y*-int was slightly below zero, driven by a few observers with larger negative *y*-int values (**Figure 6**), and did not differ between experiments (−0.50% for Discrimination, −0.54% for Estimation). Interestingly, amplitude estimates were higher for all trial events in the Discrimination experiment compared to the Estimation experiment (**Figure 7a**, *χ*^2^(1) > 32.62, p < 1.23e-7 with Bonferroni correction for multiple comparisons across parameters). To assess whether this effect was present at an individual observer level, we examined the trial event amplitudes of the three observers who participated in both experiments. We found that two out of the three observers, like the group data, had higher event amplitudes for Discrimination compared to Estimation (differences of 6.2013% and 8.4884%), whereas one had similar amplitudes (difference of −0.2353%). No other task differences survived correction for multiple comparisons.

Latency estimates were similar for the two experiments (**Figure 7b**, *χ*^2^(1) < 1.75, p > 0.18). The latency estimate for T2 was similar to the event onset, (51.4612 ms for Discrimination, 18.6015 ms for Estimation, comparison to zero latency: *χ*^2^(1) = 2.62, p = 0.11). However, the latency estimate for T1 was well before T1 onset (−290.3101 ms for Discrimination, −181.8491 ms for Estimation, comparison to zero: *χ*^2^(1) = 65.88, p = 1.92e-15 corrected). The precue latency estimate was also well before the precue onset (−156.9818 ms for Discrimination, −221.0213 ms for Estimation, comparison to zero: *χ*^2^(1) = 32.13, p = 5.76e-8 corrected). Meanwhile, the response cue latency estimate was delayed relative to the response cue onset (169.2629 ms for Discrimination, 291.8451 ms for Estimation, comparison to zero: *χ*^2^(1) = 89.84, p < 8e-16 corrected). The mean *t*_max_ was 1,053.3 ms for Discrimination and 1,295.8 for Estimation, with no significant difference (*χ*^2^(1) = 0.12, p = 0.73). Thus the two experiments had similar latency but different amplitude profiles, with larger event-related pupil responses in the Discrimination than the Estimation experiment.

## Discussion

Pupil size is an accessible, continuous measure that reflects rapidly changing internal states, but the pupil response itself is relatively slow. Linear modeling has shown promise for inferring the dynamics of internal signals that drive pupil responses, but as has been noted (Bach et al., 2018), these methods have not been systematically evaluated. To work toward a standard pupil modeling approach, here we compared different pupil models, validated modeling procedures, and evaluated the reliability of the best model. Based on the results, we recommend a specific pupil model and fitting procedure, and we quantify the uncertainty of the resulting parameter estimates. The best model includes timing parameters that are not usually fit, indicating that more precise modeling of pupil dynamics may improve the estimation of pupillary responses to rapid events.

### Model validation

Despite the increasing use of linear models of pupil size to capture pupillary time series to multiple sequential events, such methods have not been well validated. Our best model and fitting procedures performed well on simulated data, with reasonably accurate parameter recovery. The model also fit the real data well. The unknown nature of noise in the pupil data limited our ability to simulate the impact of noise on parameter recovery, so we also quantified the uncertainty of the parameter estimates in the real data. Note that these uncertainty estimates are expected to depend on the number of trials. We also identified a few parameter tradeoffs in the real data. Both uncertainty and tradeoffs should be considered when interpreting parameter estimates and potentially when designing experiments. For example, given the ∼350 ms 95% confidence interval on latency estimates, it may be helpful to separate successive trial events by at least that interval, if possible. The results suggest that the current model is reasonable as a current standard and can serve as a starting point for future work.

### Temporal properties

The inclusion of two timing parameters, event latency and *t*_max_, improved the model’s ability to fit the pupil size time series. With respect to latency, internal signals related to the precue and T1 events were estimated to occur before the events themselves. This finding suggests that these internal signals anticipated the stimulus onsets, which were predictable. Pupil dilation in advance of a predictable stimulus has been observed previously and found to depend on temporal expectation (Akdoğan, Balci, & van Rijn, 2016; Bradshaw, 1968) and upcoming task demands (Irons, Jeon, & Leber, 2017). Allowing for variable latency in pupil models may therefore be particularly important when observers have expectations about the timing of upcoming events. More broadly, latency estimates can provide information about anticipatory processes related to the observer’s task.

Most previous pupil modeling studies (de Gee et al., 2017; de Gee et al., 2014; Kang et al., 2014; Kang & Wheatley, 2015; Lempert et al., 2015; Murphy et al., 2016; van Kempen et al., 2019; Wierda et al., 2012; Willems, Damsma, et al., 2015; Willems, Herdzin, et al., 2015; Zylberberg et al., 2012) have used the canonical puRF proposed by Hoeks and Levelt (1993), which assumes that all observers have identical pupil dynamics. We found, on the contrary, that fitting the time-to-peak (*t*_max_) of the puRF improved model fits. The value of *t*_max_ varied widely across observers but was highly consistent for a given observer, suggesting that *t*_max_ is an observer-specific property. The *t*_max_ values we estimated for individual observers were in line with Hoeks and Levelt’s original estimates using a single stimulus event, which ranged from 630-1300 ms and showed some variability between auditory vs. visual events (Hoeks & Levelt, 1993). van den Brink et al. (2016) also varied *t*_max_, but did so by setting it to the latency of the maximum dilation in the time series, rather than fitting it. Here we found that individual puRFs can be estimated from a multi-event time series and should be used instead of the canonical puRF to improve pupil modeling.

Despite the sluggishness of the pupillary response, a model with two target events outperformed a model with only one target event. This suggests that the two target events were associated with separate pupil dilations, even though they were separated by only 250 ms. Due to the early response to T1, however, the estimated dilations occurred further apart in time, closer to 500 ms. Individual observer latency estimates had a reliability of ∼350 ms, indicating that while separate pupillary responses to events close in time seem to be recoverable, one should take care in interpreting their exact timing. The interpretation of some estimated latencies in the current data set was also limited by the fact that they fell at the boundary of the allowed range, −500 ms.

### Dependence on task and temporal attention

The event-related pupil response amplitude depended on the task observers were performing, with larger amplitudes in the Discrimination compared to the Estimation task. This finding demonstrates that even when the stimulus sequence is identical, cognitive factors can influence the pupil response, consistent with a large body of research (Ebitz & Moore, 2018; Einhäuser, 2016; Mathôt, 2018). Here, a relatively modest change in task instruction – discrimination vs. estimation – changed the amplitudes of pupillary responses to sensory stimuli. The larger amplitude effect for Discrimination could have been due, at least in part, to a larger baseline pupil size in the Estimation task, perhaps related to a higher cognitive load, as tonic size and phasic response amplitudes are inversely related (Aston-Jones, Rajkowski, Kubiak, & Alexinsky, 1994; de Gee et al., 2014; Gilzenrat, Nieuwenhuis, Jepma, & Cohen, 2010; Murphy, Robertson, Balsters, & O’Connell R, 2011). We were unable to directly compare baseline pupil size across tasks, however, because the tasks were performed in separate sessions. Task affected pupil response amplitudes to trial events more than the sustained, task-related amplitude, and had no effect on pupil response latencies. These results show how models can help specify the effects of cognitive manipulations on pupillary responses.

In contrast, we found no reliable impact of temporal attention on any model parameter, despite finding overall effects of temporal expectation in the form of anticipatory responses to the predictably timed stimuli, as well as behavioral effects of temporal attention in the same experiments (Denison et al., 2017). While it is difficult to draw any strong conclusions from a null result, our findings suggest that the effects of voluntary temporal attention on pupil size may be subtle, if not absent. Previous research has linked changes in pupil size to temporal selection during the attentional blink, in which observers identify targets embedded in a rapid visual sequence (Wierda et al., 2012; Willems, Damsma, et al., 2015; Willems, Herdzin, et al., 2015; Zylberberg et al., 2012). One model of the attentional blink explains the phenomenon as arising from the dynamics of the LC (Nieuwenhuis, Gilzenrat, Holmes, & Cohen, 2005), according to the idea that phasic LC responses act as a temporal filter (Aston-Jones & Cohen, 2005). Temporal selection during the attentional blink – in which target timing is unpredictable – may be different, however, from voluntary temporal attention – the prioritization of specific, relevant time points that are fully predictable. In contrast to our findings for the pupil, microsaccade dynamics are modulated by both temporal expectation (Amit, Abeles, Carrasco, & Yuval-Greenberg, 2019; Dankner, Shalev, Carrasco, & Yuval-Greenberg, 2017; Denison et al., 2019; Hafed, Lovejoy, & Krauzlis, 2011; Pastukhov & Braun, 2010) and temporal attention (Denison et al., 2019).

### Limitations and extensions

The best model was determined based on a sample dataset in which ∼2-s trials contained multiple sequential events. This dataset was therefore suited to ask about the recovery of rapid internal signals driving pupil dilation. Nevertheless, other models may perform better for different datasets. For example, Murphy et al., 2016, showed that the task-related component, which we found to be best modeled as a boxcar, was better described as boxcar or linear depending on the task. van Kempen et al., 2019, also found support for a linear task-related component. Note that all of these best-fitting boxcar and linear regressors were dependent on RT, unlike the linear regressor we tested, which modeled linear drift across the whole trial following Wierda et al., 2012.

Future work could extend the current model in multiple ways. In the main analyses, we modeled average pupil time series across trials in a given experimental condition, using the median RT to define the task-related boxcar component. Another approach would be to model single trial time series and define the boxcar for each trial using that trial’s RT (e.g., de Gee et al., 2014). In the present study, we tested only the *t*_max_ parameter of the puRF, leaving the second parameter, *n*, fixed. The motivations for this choice were that 1) *t*_max_ is easily interpretable as the time-to-peak of the puRF, whereas *n* governs the shape of the puRF in a more complex way, and 2) Hoeks and Levelt reported that *n* could vary considerably without a large impact on the other parameter estimates. However, future work could test whether fitting *n* would further improve the model fits. Future work could also add covariates to the model such as tonic pupil size (de Gee et al., 2014), model pupil modulations associated with blinks and microsaccades (Knapen et al., 2016), and model pupil constrictions as well as dilations (Korn & Bach, 2016). A recent pupil model included both transient and sustained components of the pupil dilation response (Spitschan et al., 2017), which could be compared to the unitary puRF used here and in most previous work.

### Pupil Response Estimation Toolbox (PRET)

We provide a MATLAB toolbox, PRET, that performs all the analyses reported here, including basic preprocessing (blink interpolation and baseline correction), model specification, model fitting, bootstrapping of parameter estimates, data simulation and parameter recovery, and model comparison. The toolbox is open-source and freely available on GitHub (https://github.com/jacobaparker/PRET). Importantly, PRET can be readily employed for model comparison and uncertainty estimation in other data sets to continue to work toward a field-standard pupil modeling approach.

### Open Practices Statement

The code for the pupil analyses is available at https://github.com/jacobaparker/PRET. The data for all experiments are available upon request. None of the experiments was preregistered.

## Supplementary Figures

**Figure S1.**
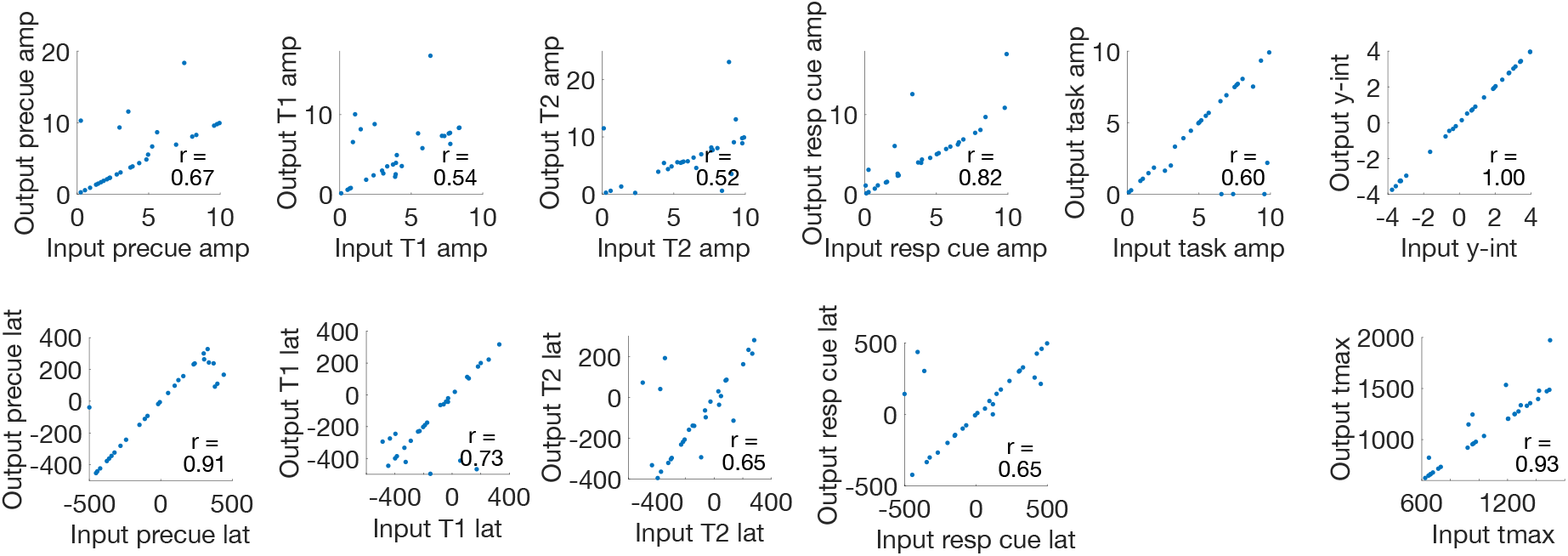
Parameter recovery for simulated data in which the actual T1-T2 SOA was 150-350 ms (i.e., +/-100 ms around the experimental T1-T2 SOA of 250 ms). All Pearson’s correlations had a corrected p-value < 0.05.

**Figure S2.**
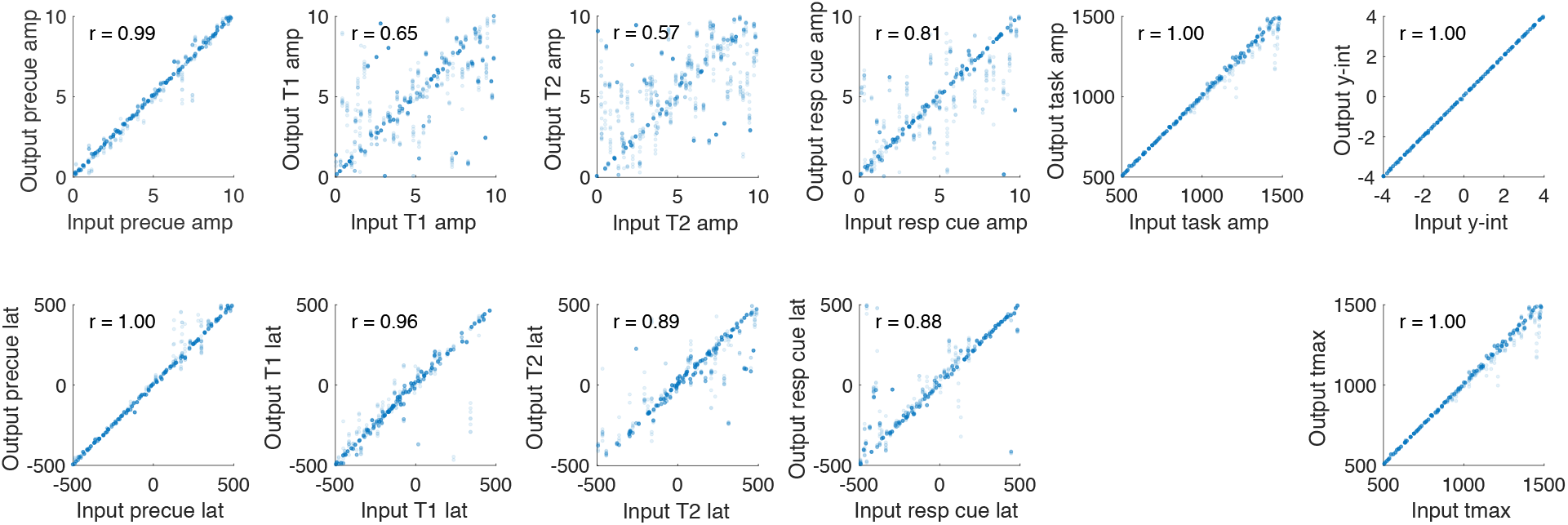
Parameter recovery for simulated single-trial data including simulated noise. Input parameters (1000 sets) for model 1 were pseudo-randomly generated from uniform distributions over the ranges shown, with binning to ensure that the whole range was sampled. During fitting, the parameters were constrained to the same ranges. Response time values used in the boxcar function for the task-related component varied from 2350 to 3350 ms. Each parameter set was used to simulate time series for 256 trials (same number of trials as in the precue T1 and precue T2 conditions of the experiment). I.i.d. noise was added to each time series, with the noise variance set to the mean squared error of the single trial fits of the real data. The median mean squared error across observers was used for the simulations. Model 1 was then fit to these simulated noisy single trials to estimate the output parameters. All Pearson’s correlations had a corrected p-value < 0.05.

**Figure S3.**
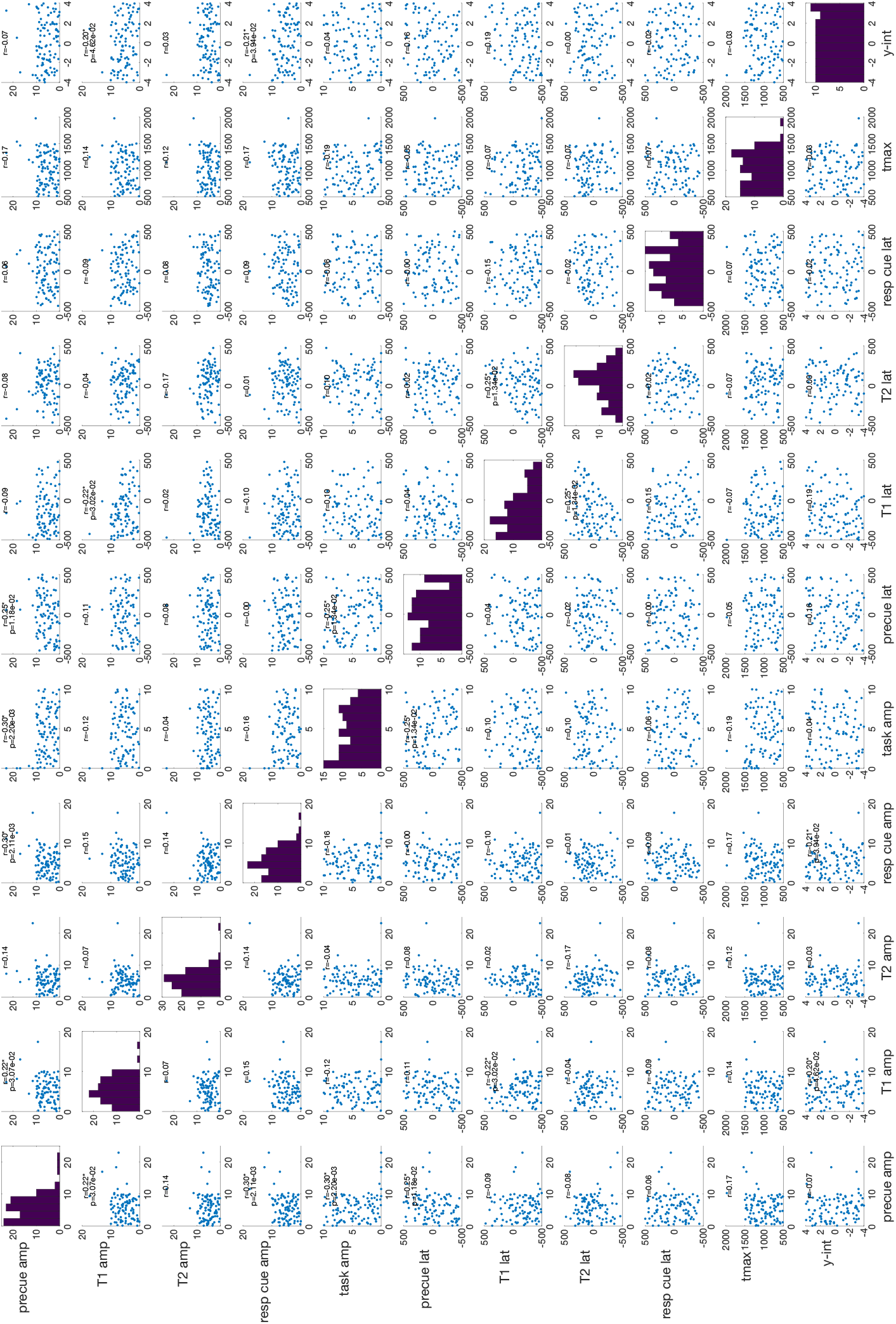
Assessing parameter tradeoffs. Scatter plots showing estimated parameter values for each parameter plotted against every other parameter estimate, from the parameter recovery simulation. Pearson’s correlation and uncorrected p-value are shown for each plot. Some correlations (9 out of 55) had an uncorrected p-value < 0.05, but none was significant after Bonferroni correction. Some of these correlations, specifically in the amplitude parameter values, were driven by the presence of a few outlier data points. Others, in the latency parameter values, were due to the non-uniform joint distributions of the parameter values caused by the temporal ordering constraint (e.g., the T2 event must follow the T1 event).

